# A large-scale dataset of functional mouse ganglion cell layer responses

**DOI:** 10.64898/2025.12.04.691221

**Authors:** Dominic Gonschorek, Jonathan Oesterle, Thomas Zenkel, Federico D’Agostino, Chenchen Cai, Nadine Dyszkant, Klaudia P. Szatko, Florentyna Deja, Tom Schwerd-Kleine, Ryan Arlinghaus, Katrin Franke, Zhijian Zhao, Timm Schubert, Philipp Berens, Thomas Euler

**Affiliations:** Institute of Ophthalmic Research, University of Tübingen, Tübingen, Germany; Werner Reichardt Centre for Integrative Neuroscience, University of Tübingen, Tübingen, Germany; Hertie Institute for AI in Brain Health, University of Tübingen, Tübingen, Germany; Tübingen AI Center, University of Tübingen, Tübingen, Germany; Department of Ophthalmology, Byers Eye Institute, Stanford University School of Medicine, USA; Stanford Bio-X, Stanford University, USA; Wu Tsai Neurosciences Institute, Stanford University, USA

**Keywords:** retina, ganglion cells, displaced amacrine cells, large-scale dataset

## Abstract

We present the All-GCL dataset, a large-scale resource of functional two-photon Ca^2+^-imaging recordings with rich meta-data information from more than 80,000 cells in the ganglion cell layer (GCL) of the *ex vivo* mouse retina. Collected over nine years across more than 139 experimental sessions, the dataset provides recordings of light-evoked responses to various stimuli, including a shared set of core stimuli. To enable cell-type–specific analyses, cells are probabilistically assigned to 46 previously characterized functional groups, including retinal ganglion cells and displaced amacrine cells. Further, we assessed the influence of experimental and biological factors on the functional responses and found only small batch effects across experimenters, setups, and recording sessions, highlighting the dataset’s consistency. The All-GCL dataset offers a comprehensive and standardised reference for studying retinal computation at scale. It supports population-level analyses, computational modelling, and the development of machine learning approaches for biological time-series data. Future releases will expand the dataset with additional mouse lines and light stimuli, creating a growing resource for the vision science community.

## Background & Summary

Sensory systems are essential for survival, enabling organisms to perceive, interpret, and respond to their environment. Among mammals, vision is often an important if not the dominant sense, supporting critical behaviours such as navigation, foraging, and social interaction. Visual perception begins when photons reach the retina – a thin, multi-layered neuronal tissue at the back of the eye. Photoreceptors within the retina detect light and transduce it into electrical signals, which are relayed through bipolar cells (BCs) to retinal ganglion cells (RGCs), the output neurons of the retina to the rest of the brain (1, 2).

The retina’s primary task is to transform the continuous flow of light patterns from the environment into parallel representations of visual features such as intensity, contrast, “colour”, and motion (3). This transformation is mediated by its highly organized circuit architecture, which integrates feedforward and lateral interactions across multiple cell classes. Amacrine cells (ACs), for instance, modulate temporal and spatial aspects of RGC responses, shaping computations such as motion detection and contrast adaptation before signals are transmitted to the brain (4, 5). Through this sophisticated processing, the retina operates not merely as a camera sensor but as a dynamic computational network that actively extracts essential, behaviourally relevant features from the visual world (6). Here, RGCs represent the retina’s output channels, encoding specific visual features and transmitting them to distinct downstream brain regions.

In the mouse retina, up to 45 functional RGC types have been identified (7, 8), a diversity mirrored by morphological and molecular classifications (9, 10). Each RGC type contributes to diverse aspects of visual encoding, ranging from edge detection and direction selectivity to luminance and chromatic contrast, also underscoring the retina’s capacity for distributed computation. Understanding how these diverse cell types collectively represent visual information remains a central question in vision research.

Progress in addressing this question increasingly depends on access to large, standardised datasets that capture the retina’s activity. While there is a plethora of studies that have provided valuable and detailed insights into individual RGC types and their local circuits, large-scale datasets that combine broad sampling with consistent experimental conditions and well-defined stimuli are still rare, or not easily publicly accessible. Yet, such resources are crucial for uncovering, for instance, response variability within RGC types, population-level principles, validating computational models, and enabling cross-study comparisons. Moreover, the increasing role of machine learning and data-driven modelling in neuroscience amplifies the need for open, well-curated, and highquality datasets that bridge biological experiments and computational analysis.

Here, we present the All-GCL dataset, a large-scale collection of functional recordings from over 80,000 cells in the ganglion cell layer (GCL) of the *ex vivo* mouse retina, obtained using two-photon Ca^2+^-imaging under standardised conditions. This dataset integrates recordings from multiple experimental studies spanning nine years, all performed with a shared battery of “fingerprinting” visual stimuli. In the following, we describe the dataset in detail, summarise its composition and data quality, quantify batch effects, and demonstrate its potential for functional classification and comparative analyses.

The overarching goal of this work is to provide a rich and accessible resource for the vision science community. The All-GCL dataset enables large-scale analyses of retinal function, supports benchmarking of computational models, and may serve as a dataset for machine learning applications involving biological time-series data. We will continue to expand this resource with additional mouse lines (e.g., retinal degeneration mutants, such as *rd10*) and new stimulus protocols, creating a reference dataset for the functional organization of the mouse GCL.

## Methods

### Input data

With the All-GCL dataset, we combine data recorded over a period of nine years. In total, the dataset contains 139 experimental sessions that were all originally collected for different studies:

- a study on colour vision by Szatko et al. (11),
- a machine-learning driven study on processing of natural scenes of RGCs by Höfling et al. (12),
- a study on the effects of nitric oxide on RGCs by Gonschorek et al. (13),
- a study on the functional degeneration of rd10 mice where they used wild-type mice as a control group by Dyszkant et al. (14),
- unpublished work by Franke et al. (2018) on retinal processing of natural scenes,
- unpublished work by Cai et al. (2022-2024) on object-background segregation in the early visual system that was already presented at conferences (15),
- and unpublished work by Kokat, Deja et al. (2022-2023) on suppressed-by-contrast RGCs in the early visual system that was already presented at conferences.

All data included share a standardised experimental procedure that features a set of “fingerprinting” stimuli, namely a full-field “chirp” and a moving bar, established by Baden et al. (7). This large-scale aggregation enables systematic comparison of light-evoked activity across preparations and conditions, and represents a comprehensive resource for studying response diversity in the mouse GCL.

### Animals and preparation

All animal experiments were conducted at the University of Tübingen and were performed according to the laws governing animal experimentation issued by the German Government as well as approved by the institutional animal welfare committee of the University of Tübingen (31.10.2016, CIN08/19M, CIN03/21M, CIN06/21M). For the pooled dataset, we used retinae (n=203) from C57Bl/6J mice (n=139; JAX 000664, The Jackson Laboratory) of either sex between the age of 3 to 30 weeks. All animals were kept in the local animal facility and housed under the standard 12h/12h day/night cycle at 22°C and a humidity of 55%.

The following procedures were carried out under very dim red (> 650 nm) light. Before each imaging experiment, the animal was dark-adapted for >1h, then anesthetized with isoflurane (CP-Pharma) and sacrificed by cervical dislocation. Immediately after, the eyes were enucleated with a dorsal cut as orientation landmark and hemisected in carboxygenated (95% O_2_, 5% CO_2_) artificial cerebrospinal fluid (ACSF) solution containing (in mM): 125 NaCl, 2.5 KCl, 2 CaCl_2_, 1 MgCl_2_, 1.25 NaH_2_PO_4_, 26 NaHCO_3_, 20 glucose, and 0.5 L-glutamine at pH 7.4. Sulforhodamine-101 (SR101, 0.1 µM; Invitrogen) was added to the ACSF to reveal blood vessels and damaged ganglion cell layer (GCL) cells in the red fluorescence channel (16). The carboxygenated ACSF was constantly perfused through the recording chamber at 4 ml/min and kept at ∼ 36°C throughout the entire experiment.

After the dissection, retinae were bulk-electroporated with the synthetic fluorescent calcium indicator Oregon-Green 488 BAPTA-1 (OGB-1; hexapotassium salt; Life Technologies) (17). To electroporate the GCL, the dissected retina was flat-mounted with the GCL facing up onto an AnodiscTM (#13, 0.1 µm pore size, 13 mm diameter, Cytiva), and then placed between two 4 mm horizontal platinum disk electrodes (CUY700P4E/L, Nepagene/Xceltis). The lower electrode was covered with 15 µl of ACSF, while a 10 µl drop of 5 mM OGB-1 dissolved in ACSF was suspended from the upper electrode and lowered onto the retina. Then, nine electrical pulses (∼ 9.2 V, 100 ms pulse width, at 1 Hz) from a pulse generator/wide-band amplifier combination (TGP110 and WA301, Thurlby handar/Farnell) were applied and then, the electroporated retina on the Anodisc was transferred into the recording chamber, whereby the dorsal edge of the retina pointed away from the experimenter. The retina was left there for ∼30 min to recover, as well as adapted to the light stimulation by displaying a binary dense noise stimulus (20 × 15 matrix, 40 × 40 µm^2^ pixels, balanced random sequence) at 5 Hz before the recordings started.

### Two-photon Ca^2+^-imaging

For the functional Ca^2+^-imaging experiments, a MOM-type two-photon microscope (Sutter Instruments/Science Products) was employed (16, 18). The microscope was equipped with a mode-locked Ti:Sapphire laser (MaiTai-HP DeepSee, Newport SpectraPhysics) tuned to 927 nm (to optimally excite OGB-1), two photomultipliers serving as fluorescence detection channels for OGB-1 (HQ 510/84, AHF/Chroma) and SR101 (HQ 630/60, AHF), and a water immersion objective (CF175 LWD×16/0.8W, DIC N2, Nikon, Germany). To acquire images, custom-made software (ScanM by M. Müller and T. Euler) running under IGOR Pro for Microsoft Windows (Wavemetrics) was used and time-lapsed 64 × 64 pixel image scans (100 × 100 µm) at 7.8125 Hz were taken. Routinely, the optic nerve position and the scan field position were recorded to reconstruct their retinal positions. High-resolution images (512 × 512 pixel images) were recorded to support region of interest (ROI) detection.

### Light stimulation

For the light stimulation of the retinal tissue, a digital light processing (DLP) projector (lightcrafter (LCr), DPM-E4500UVBGMKII, EKB Technologies Ltd) was used to display the visual stimuli through the objective onto the retina, whereby the stimulus was focused on the photoreceptor layer (19). To enable differentially exciting the two spectral mouse cone types (PR1 and PR4, with peak sensitivities of 510 and 360 nm, respectively), the stimulator used two narrow bands at around ∼ 575 nm and ∼385 nm. This was implemented either by equipping the LCr itself with the respective LEDs and band-passing the projected image using a dual-band filter (390/576 Dualband, F59-003, AHF/Chroma) or by using a light-guide enabled LCr with an external, individually band-pass filtered green/UV LED unit (LEDs; green: 576 BP 10, F37-576; UV: 387 BP 11, F39-387; both AHF/Chroma; for details, see ref. (19)).

In either configuration, the LEDs were synchronized with the scan retracing of the microscope. Stimulator intensity (as photoisomerization rate, 10^3^ P^∗^s^−1^ per cone) was calibrated to range from ∼0.5 (“black” image) to ∼20 for M-(LWS) and S-(SWS1) opsins, respectively. Steady illumination of ∼10^4^ P^∗^s^−1^ per cone was present during the scan recordings due to the two-photon excitation of photopigments (16, 18).

Two types of light stimuli were used in all experimental sessions included in this dataset: a full-field “chirp” stimulus (with either 800 µm or 1000 µm ∅; see details in ref. (7)), and a bright moving bar (0.3 × 1 mm) at 1 mm s^−1^ in eight directions to probe direction and orientation selectivity. Additionally, for a subset of the experiments, a balanced random binary noise stimulus with a checkerboard grid of 20 × 15 checks and a check size of 40 µm at 5 Hz was displayed for 5 min to map receptive fields.

Light stimulus centre and scan field centre were aligned for all stimuli. Before each stimulus was presented, the baseline was recorded after the laser started scanning for at least 10 s to allow any immediate laser-induced effect on the retinal activity to have subsided (11, 16, 18).

## Data analysis

### Software

Image extraction was performed using IGOR Pro. All analyses were performed in Python and the results were stored and organised in a custom DataJoint database (20).

### Regions of interest

Regions of interest (ROIs), which correspond to the somata of individual cells in the GCL, were manually drawn in the input data using IGOR Pro. Some of the input data originally excluded cells with strong SR101 filling from the ROI masks because it was assumed that SR101 filling correlated with cell death. Here, however, we wanted to include all cells in the ROI masks. To this end, we added previously unlabelled cells to the ROI masks as follows: For each field, we generated a new temporary ROI mask using Cellpose 3 (21). If any new ROIs were missing from the manually drawn ROI masks (defined as having a maximum intersection-over-union of 0.1 with any other ROI), we added them to the manual ROI mask after clipping any overlapping or directly adjacent pixels.

### Preprocessing

Ca^2+^-traces were extracted from individual ROIs as the mean over all pixels in the ROI for each imaging frame. These raw traces were detrended by subtracting a smoothed version *r*_*smooth*_ of the trace from the raw one. Detrending was necessary to remove slow drifts in the signal that were unrelated to the light-induced response. The smoothed trace *r*_*smooth*_ was computed by applying a Savitzky-Golay filter (22) of 3^*rd*^ polynomial order and a window length of 60 s using the Python SciPy implementation (23).

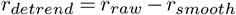

For the “chirp” and moving bar stimulus, we further computed response averages from the detrended traces as means over repetitions. In the case of the moving bar stimulus, the response average was reduced to the response to the preferred motion direction of the cell (for details, see ref. (7)). Finally, response averages were normalised by first subtracting the baseline activity (computed as the mean over the first second), and then by dividing by the maximum amplitude *max*_*t*_( |*r*(*t*) |) = 1. This normalization was performed independently for each ROI, stimulus, and condition. Direction and orientation selectivity – qualified as DSi and OSi, respectively – were computed from the moving bar responses using a permutation test as described in Baden et al. (7). The On-Off index (OOi) describes the response polarity preference of a cell to either light increments and/or decrements, and was computed as described in Baden et al. (7).

### Response quality

To quantify the response quality of individual ROIs, we computed a quality index (QI) for the moving bar (QI_MB_) and full-field “chirp” (QI_chirp_):

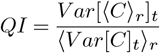

where *C* is the *T* by *R* response matrix (time samples by stimulus repetitions) and ⟨⟩_*x*_ and *V ar*[]_*x*_ denote the mean and variance across the indicated dimension *x*, respectively. For the moving bar, the QI was computed for all direction independently using the direction with the largest QI as QI_MB_. Cells with either QI_MB_> 0.6 or QI_chirp_> 0.35 were considered *responsive* in the following analysis. A field is considered responsive if at least a third of the ROIs have a QI_MB_> 0.6.

### Classification of functional ganglion cell layer groups

For the functional classification of GCL types, we built on apreviously published GCL classifier (13, 24), which is a *scikit* − *learn* random forest classifier trained on previously published GCL responses and functional cell group labels (7). As in the previous version, the classifier predicts a functional group label for each cell independently using the following inputs:

- 20 features from the normalized response average to the full-field “chirp”,
- 8 features from the normalized response average to the moving bar projected to the preferred direction,
- a p-value from a permutation test for direction selectivity,
- and the ROI sizes as a proxy for soma-size.

The features of the “chirp” and moving bar responses were extracted using the sparse PCA embedding matrix that was used in Baden et al. (7) for clustering. We split the labelled training data (n=7,979 ROIs) into a train (90%) and held-out test (10%) set using stratification. We used the original 75 cluster labels from Baden et al. (7) as the classifier output. Hyperparameters were tuned using a cross-validated grid search. Using the final set of hyperparameters, the random forest classifier underwent uncertainty calibration (using 5-fold cross-validated probability calibration with logistic regression) and was then evaluated on the held-out test-set achieving an accuracy of 0.764 (at a chance level of 0.013). Finally, using the same hyperparameters, the classifier was re-trained on the full Baden et al. (7) dataset, i.e. including the test data.

As this model will now be used solely for prediction rather than validation, merging the test set into the training data ensures the classifier learns from the full distribution of the original study. The classifier outputs a probability distribution over the 75 cluster labels. In a post-processing step, we merged the 75 clusters to the 46 functional groups G_1_ to G_46_ as in Baden et al. (7), summing the respective classifier output probabilities. A cell is assigned to the group for which the classifier predicts the largest probability; we define this largest probability as the classification confidence of the uncertainty-calibrated classifier. To predict group labels for new data, we replicated the preprocessing of Baden et al. (7) as much as possible and applied it to the new data. However, the “chirp” they used was slightly different in its timing properties to the “chirp” used in this dataset. To align the responses as good a possible, we warped the time of the new data to maximize agreement with the Baden et al. (7) dataset.

#### Receptive field estimation

We mapped receptive fields (RFs) of cells using the RF Python toolbox *RFEst* (25), following the procedure in (7) with few modifications. The binary dense noise stimulus (20 × 15 matrix, (40 µm)^2^ pixels, balanced random sequence; 5 Hz) was centred on the recording field. We computed the temporal gradients of the Ca^2+^-signals from the detrended traces and clipped negative values:

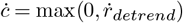

The stimulus *S*(*t*) and the clipped temporal gradients *c* were upsampled to 10 times the stimulus frequency to compute the gradient-triggered average stimulus:

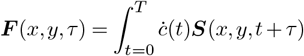

where ***S***(*x, y, t*) is the stimulus, *τ* is the lag ranging from approximately 0.20 to 1.38 seconds, and *T* is the duration of the stimulus. We smoothed these raw RFs using a 3 × 3 pixel and 1 pixel standard deviation Gaussian window for each lag. Then we decomposed the RF into a temporal (*F*_*t*_(*τ* )) and spatial (***F*** _*s*_(*x, y*)) component using singular value decomposition and scaled them such that max(|*F*_*t*_|) = 1 and max(|***F***_*s*_ |) = max(|***F*** |). RF quality was computed as:

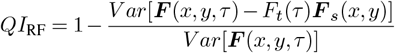

Only RFs with *QI*_RF_ > 0.4 were used for the analysis.

For each spatial RF ***F***_*s*_, we fit a 2D Gaussian using the Python package *astropy* (26). The area covered by two standard deviations of this Gaussian fit was used as the RF size.

### Estimating batch effects

To separate biological signals from potential effects induced by confounding variables in the dataset – so-called batch effects – we trained classifiers to predict metadata attributes, such as the animal’s sex or the experimenter, solely from the recorded neural activity.

We first filtered the dataset by selecting traces with the following response criteria: QI_chirp_>0.35 or QI_MB_>0.6. Based on a random seed, the filtered dataset was split into training (80% of cells) and test (20% of cells) sets based on recording dates. This ensured that the test set evaluated performance on previously unseen dates, preventing data leakage. To avoid class imbalance, we downsampled the majority class to ensure a random classifier would achieve an average accuracy of 50% assuming two classes.

To create the input features of the classifier, we concatenated the preprocessed “chirp” (259 time points) and moving bar responses to the preferred direction (32 time points) into a 291-dimensional feature vector. Then we began by training a model to distinguish between the three recording setups. Because distinct setups proved identifiable, we restricted all subsequent experiments to setup ID 1, the setup used for most recordings. For all remaining predictions, we binarised the metadata attributes for a binary classification task. Data points falling between binary thresholds were excluded to ensure separation. We trained classifiers to predict the sex of the animal (female/male), scan field ID (first/third field), retinal location (ventral/dorsal relative to the optic nerve, excluding cells within 100 *µm* of the optic nerve), and age (younger than 10 weeks & older than 15 weeks). Finally, to assess experimenter effects, we trained models to differentiate between pairs of experimenters.

For all tasks, we use a high-capacity random forest classifier (1,000 estimators, maximum depth of 50). To ensure robustness, we performed 20 independent training runs with different random seeds for the train/test split.

### Topographic mapping

To compute cell-type specific spatial density maps, we restricted the dataset to responsive cells with information about their location. Next, we computed a 2D Gaussian kernel-density estimate (KDE) for each celltype *ct* independently *d*_ct_(*x, y*). The KDE were evaluated on a regular grid of 51 × 51 points spanning 2 mm from the optic disc centre. We normalized the cell-type specific estimates using a KDE map over all cell types:

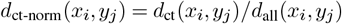

for all *x*_*i*_ and *y*_*j*_ on the grid.

## Data Record

The complete dataset generated in this study is publicly available on the Hugging Face Hub from the repository huggingface.co/datasets/eulerlab/all-gcl. Here, the data are standardised according to the Neurodata Without Borders (NWB; (27)) specifications to ensure inter-operability and long-term accessibility.

The dataset is organized hierarchically by experimental session. The root directory contains folders for each session (naming convention: `Session_Experimenter_ID_Date`), which branch into subfolders for specific experiments and anatomical locations (e.g., `experiment_1_LR` for **l**eft **r**etina). Inside these subdirectories, individual . `nwb` files represent distinct recording fields (e.g., `field_GCL1.nwb`). Additionally, a single aggregated summary table (`all_GCL_table.parquet`) is provided at the root level, containing extracted features for all cells across all experiments to facilitate rapid analysis without parsing individual . `nwb` files.

Each . `nwb` file includes both the raw Ca^2+^-imaging data and the processed results. The internal structure follows the NWB standard, with raw acquisition data stored under `acquisition/` (organized by stimulus condition: fullfield “chirp”, moving bar, dense noise) and processed results under `processing/`. Key metadata and analysis outputs include:

- *General metadata*. Experimental details including animal age, sex, genetic line, eye orientation, and anatomical position (ventral-dorsal/temporal-nasal coordinates relative to the optic disc).
- *Functional classification*. Functional cell group labels (cluster IDs 1–75), confidence scores, and probability distributions aligned with the Baden et al. (7) classification scheme.
- *Stimulus responses*. Preprocessed response traces and quality indices (Signal-to-Noise Ratio) for “chirp” and moving bar stimuli.
- *Feature indices*. Computed functional metrics, including DSi, OSi, and OOi (see above).
- *Receptive fields*. Spatio-temporal RFs derived from dense noise stimuli, including spatial parameters (RF diameter, Gaussian fit quality) and temporal dynamics (lag, transience).

Detailed descriptions of all variable names, column headers, and unit specifications are provided in the README.md file accompanying the data repository.

## Data Overview

In total, the All-GCL dataset contains responses from 84,451 GCL somata, with ∼85.5% (72,475 cells) having information about their exact retinal recording location (Fig. 2a,e). The spatial distribution of recordings covers almost the entire retinal surface, with a slight sampling bias toward ventral regions (Fig. 2e). To ensure data quality, responses were evaluated using a quantitative quality index (QI), with ∼65.6% (52,189 cells) exceeding the selected response criterion (QI_chirp_ ≥ 0.35 or QI_MB_ ≥ 0.6; Fig. 2g,h; see Methods for details).

**Fig. 1.**
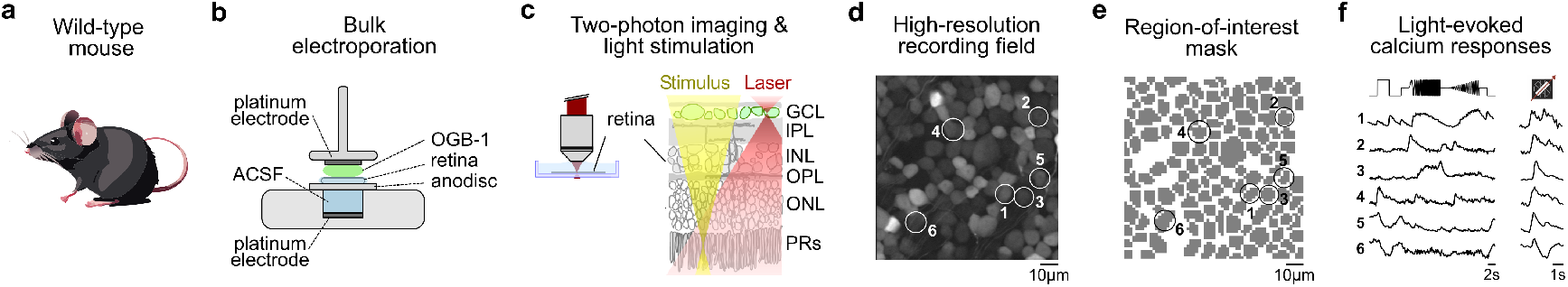
Data acquisition. (**a**) Illustration of a wild-type mice with the genetic background *C57BL/6J*. (**b**) Schematic of the procedure to bulk electroporate a whole-mounted *ex vivo* retina. (**c**) Two-photon imaging of ganglion cell layer (GCL) somata in the whole-mounted *ex vivo* mouse retina. (**d**) Representative high-resolution recording field showing GCL somata loaded with Ca^2+^-indicator Oregon Green BAPTA-1 (OGB-1). (**e**) Semi-automatic computed region-of-interest (ROI) masked drawn on the 64 × 64 pixel scan field from the highresolution field in (d) to extract individual cell responses. (**f**) Representative Ca^2+^-activity from cells in the GCL (white & black circles in (d) & (e)) in response to full-field “chirp” (left) and moving bar stimulus (right).

**Fig. 2.**
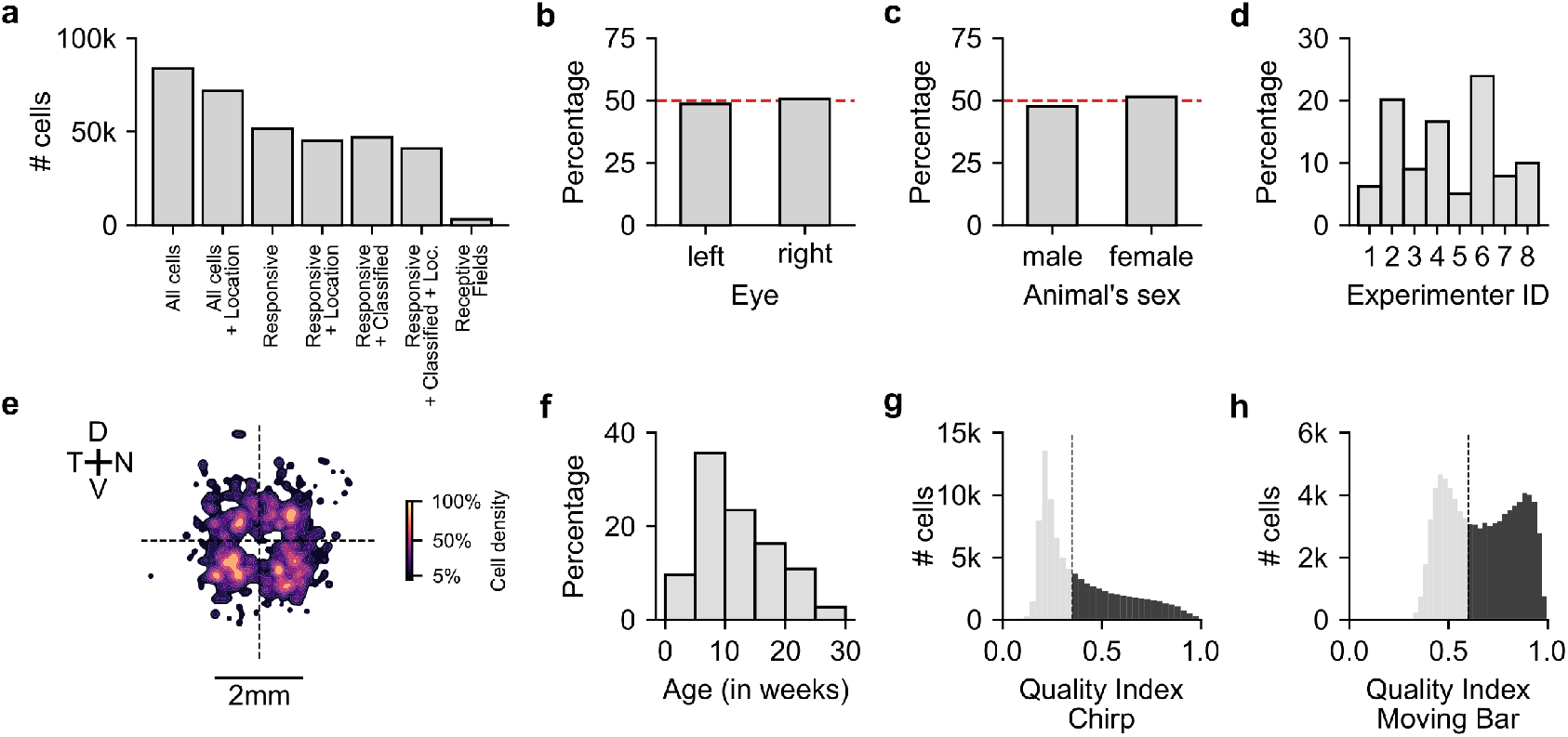
Data overview. (**a**) Number of recorded cells across different selection and response criteria. (**b-c**) Distribution of recordings across eyes (left vs. right) and animal sex (male vs. female), respectively. (**d**) Proportion of cells recorded by each experimenter. (**e**) Spatial distribution of recorded cells within the retina, shown as a density map (D, dorsal; V, ventral; T, temporal; N, nasal); Scale bar: 2 mm. (**f**) Age distribution of animals used for all recordings. (**g-h**) Distribution of response quality indices for full-field “chirp” (g) and moving bar (h) stimuli, with dashed lines indicating quality thresholds.

The dataset is well balanced with respect to eye location and sex: approximately half of the cells were recorded from left eyes and the other half from right eyes (Fig. 2b), and recordings are similarly split between male and female animals (Fig. 2c). Recordings were contributed by eight experimenters (Fig. 2d). The animals used for recordings spanned a broad age range from postnatal day (P) 15 to 200, with the majority between 3 and 30 weeks of age (Fig. 2f), thus encompassing both juvenile and mature animals.

Together, these data provide a large-scale, well-balanced, and high-quality foundation for type-specific analyses of light-evoked GCL activity. The scale and consistency of the All-GCL dataset enable reliable cross-comparisons across experiments, forming the basis for subsequent analyses.

### Functional GCL group classification

To understand how the retina segregates visual information into parallel output channels, one can identify cell groups within the GCL sharing functional response features. Using this approach, Baden et al. (7) showed that the GCL comprises approximately 46 functional groups, including 32 retinal ganglion cell (RGC) groups and 14 displaced amacrine cell (dAC) groups. Building on this classification framework, we sought to extend group-specific classification to the All-GCL dataset to enable systematic, large-scale analyses of functional diversity across recordings.

For this purpose, we employed an updated version of the previously published GCL-group classifier ((24); for details, see Methods). In brief, the supervised classifier was trained on features extracted from full-field “chirp” and moving bar responses, complemented by soma size and direction-selectivity parameters from Baden et al. (7). The group labels from that study served as ground-truth group labels. To predict cell groups in the All-GCL dataset, each recorded response was projected into the same feature space, and the classifier assigned a predicted group label along with an associated confidence score. Depending on the desired analysis level, each cell is annotated with three hierarchical labels: (1) functional group, (2) functional supergroup (i.e., ‘Off’, ‘On-Off’, ‘Fast On, ‘Slow On’, ‘Uncertain RGC’, ‘dAC’), and (3) GCL class (RGC, dAC), each accompanied by its corresponding confidence score.

Applying this refined classifier to the All-GCL dataset yielded group predictions for all recorded cells, spanning the full range of known functional groups (Fig. 3a-c). The resulting population structure closely reflects the distribution reported by Baden et al. (7), with comparable proportions of ‘Off’, ‘On–Off’, ‘Fast On’, ‘Slow On’, ‘Uncertain RGCs’, as well as ‘dACs’ (Fig. 3c). For each group, we also computed the average cell count per field with certain cells being more frequent than others (Fig. 3d). Representative average responses to the “chirp” and moving bar stimuli for selected groups illustrate the diversity of temporal and directional tuning among identified GCL types (Fig. 3b,e).

**Fig. 3.**
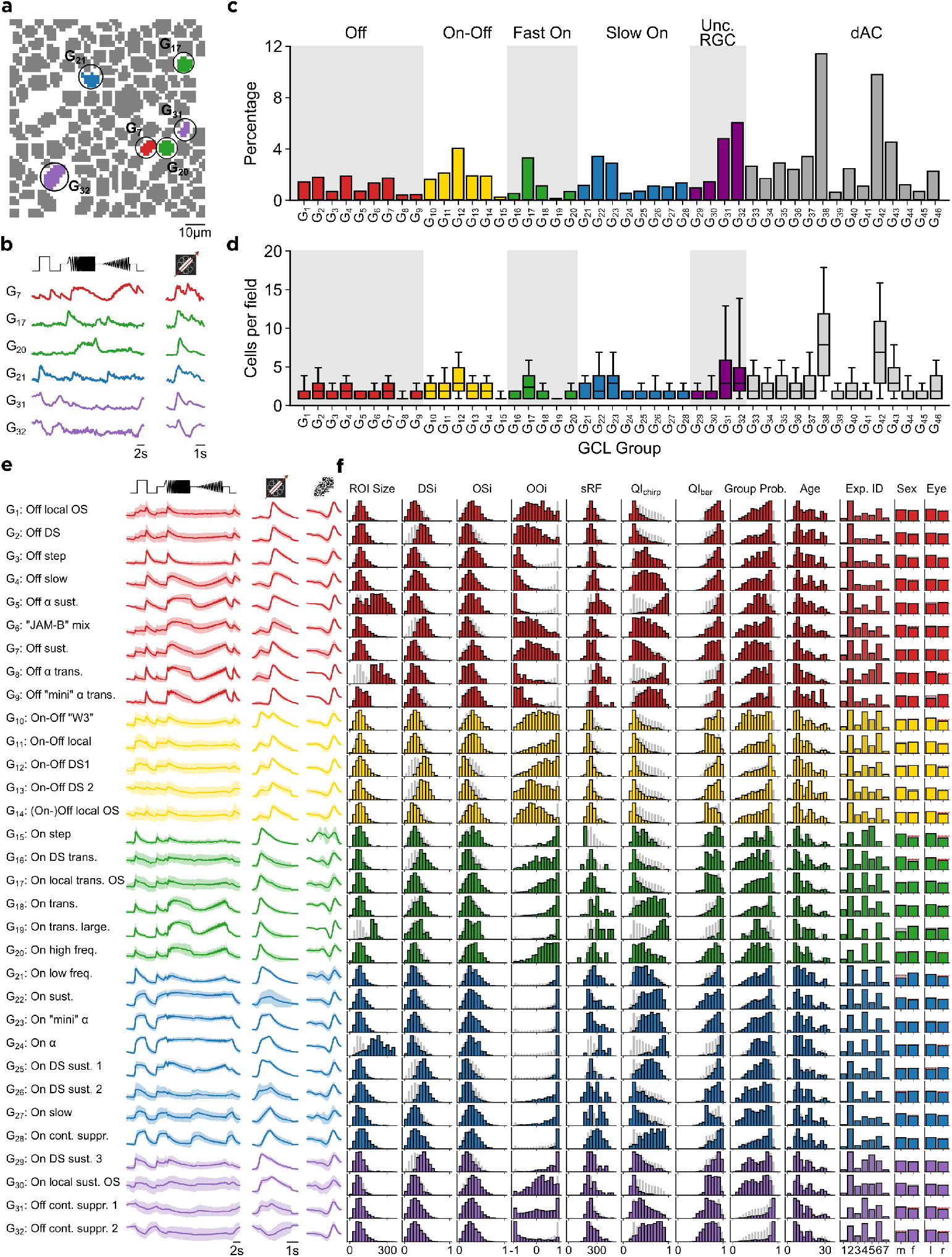
Functional classification of GCL somata. (**a**) Semi-automatic drawn region-of-interest (ROI) mask on top of the recording field from Fig. 1a with examples of functionally classified ganglion cell layer (GCL) groups. (**b**) Representative average “chirp” and moving bar responses from the recording field in Fig. 1a,b, colour-coded according to their classified functional group. (**c**) Distribution of cell groups across functional supergroups, separated into ‘Off’, ‘On-Off’, ‘Fast On’, ‘Slow On’, ‘Uncertain RGCs’, and displaced amacrine cells (dACs). (**d**) Average number of cells per recording field for each functional group. (**e**) Mean “chirp”, moving bar responses, and temporal receptive field for all identified GCL groups, organized by functional group and colour-coded consistently with panels b-d. (**f**) Corresponding distribution of key physiological and experimental parameters for each group, including ROI size, direction selectivity index (DSi), orientation selectivity index (OSi), On-Off index (OOi), spatial receptive field area (sRF), “chirp” and moving bar response quality indices, and metadata (age, experimenter ID, sex, eye).

To further characterize each group, we computed and added the distributions of key physiological and morphological parameters, including ROI size (reflecting soma size), DSi, OSi, OOi, spatial RF size, quality indices for “chirp” and moving bar, probability of the group label classification, age range, experimenter ID, animal’s sex, and eye (Fig. 3f). Across groups, these features followed characteristic distributions that matched previously described response motifs. The consistent mapping of these properties validates the reliability of the classifier across the full dataset.

Importantly, the predicted group, supergroup, and class labels, as well as the corresponding confidence scores are made available as part of the All-GCL dataset. These labels provide users with an immediately usable framework for querying, filtering, and analysing GCL responses by functional group. Together, they enrich the dataset with biologically meaningful structure and enable researchers to perform group-specific analyses without additional preprocessing.

### Technical Validation

#### Validating RGC groups through their topographical distributions

Previous studies have characterized topographic distributions of RGC types using spatial transcriptomics (e.g., (9, 28, 29)), immunohistochemistry/transgenic lines (e.g., (30–32)), and functional recordings for a subset of types (e.g., (7, 33, 34)). For several groups, particularly alpha RGCs (35), functional identities align well with molecular and anatomical descriptions (8). These prior findings provide a reference for validating the spatial organization present in the All-GCL dataset.

To assess consistency with earlier studies, we computed spatial density maps for all 46 functional GCL groups (Fig. 4), expressed as relative density normalized with respect to the density of all cells. At the level of functional supergroups, the observed distributions agreed with established large-scale patterns: ‘Off’ groups (G_1_–G_9_) showed dorsal enrichment, while ‘Fast On’ and ‘Slow On’ groups (G_15_–G_19_ and G_20_–G_28_) were enriched ventrally. ‘On–Off’ groups (G_10_–G_14_), as well as ‘Uncertain RGCs’ (G_29_–G_32_) and ‘dACs’ (G_33_–G_46_), exhibited weaker or more heterogeneous gradients, consistent with earlier anatomical and functional observations.

**Fig. 4.**
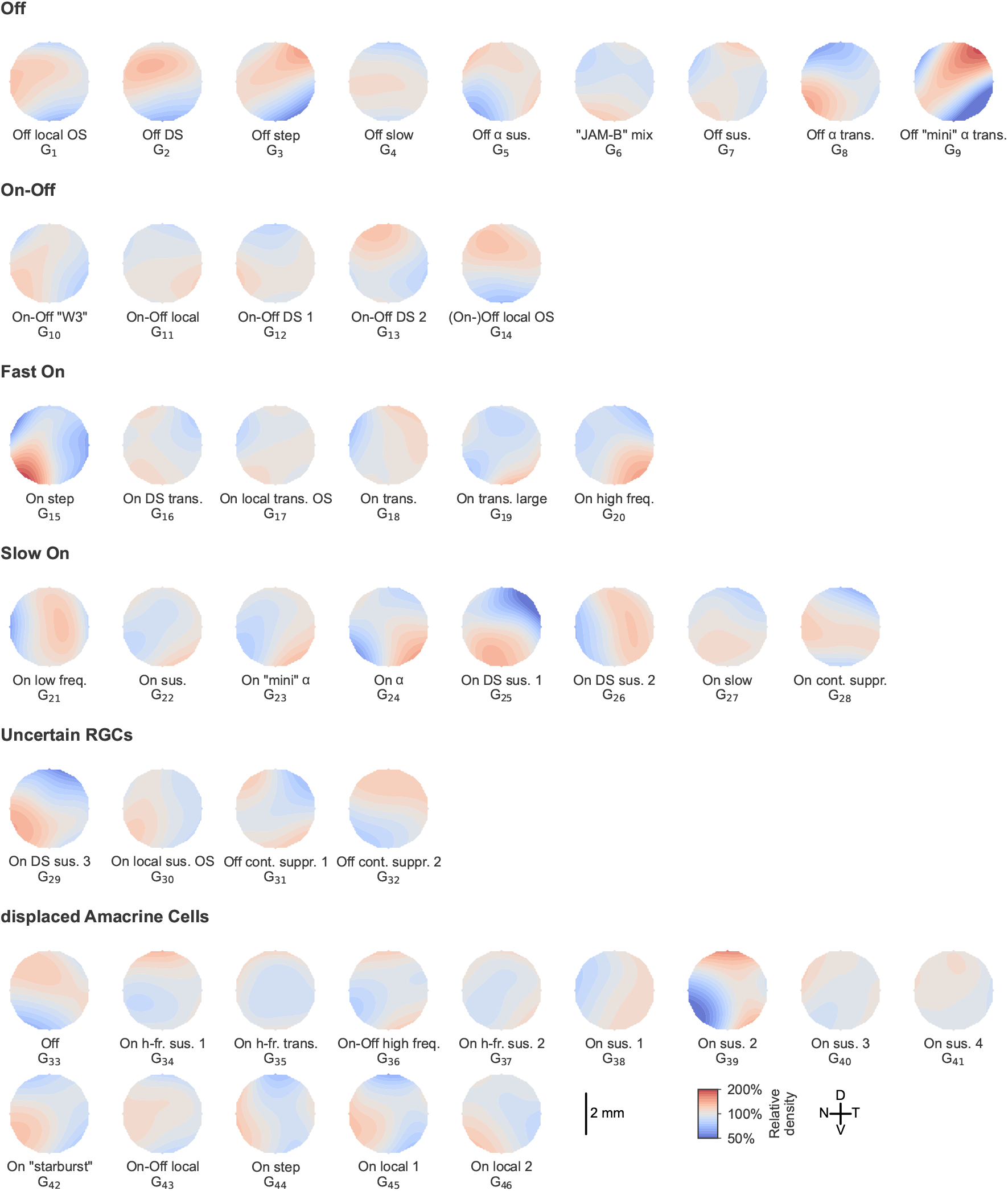
Topographic distribution of functional GCL groups. Spatial distribution maps of all identified functional GCL groups across the retina. Each circle represents the relative density of a specific functional group, with colour indicating the normalised cell density (red: higher density, blue: lower density). Functional groups are arranged by response class: ‘Off’, ‘On–Off’, ‘Fast On’, ‘Slow On’, ‘Uncertain RGCs’, and ‘displaced amacrine cells’. D, dorsal; V, ventral; T, temporal; N, nasal. Scale bar, 2 mm.

For individual groups with well-documented topography, the maps reproduced expected trends. The On alpha sustained group (G_24_) showed temporal-ventral enrichment in agreement with previous studies (28–31, 36), and the On contrast suppressed group (G_28_) showed a naso–temporal bias consistent with recent functional findings (12). The overall complementarity of dorsal ‘Off’ and ventral ‘On’ distributions matches established retinal specializations (11, 37).

Together, these correspondences demonstrate that the spatial organization of functional groups in the All-GCL dataset aligns well with published molecular, anatomical, and functional topographies, providing technical validation for the group assignments.

#### Quantifying batch effects in the dataset

Variability introduced by extraneous, non-biological factors – commonly referred to as batch effects – can obscure or confound meaningful biological signals in datasets. Such effects may arise from subtle differences in experimental conditions, handling procedures, or systemic variation across data collection sessions. For example, Zhao et al. (38) observed subtle batch effects in retinal data, namely changes in the neural response kinetics, likely induced by temperature fluctuations during the recordings. To assess the extent of such influences in the All-GCL dataset, we examined six key metadata parameters: recording setup ID, experimenter identity, scan field ID (first or third recording field), retinal location (ventral or dorsal), animal’s sex (male or female), and age (juvenile or adult). Among these, sex, age, and retinal location represent biological covariates that could reflect intrinsic physiological variation, or ecological adaptations, whereas setup ID and experimenter identity constitute technical covariates, capturing procedural inconsistencies across recordings. The scan field parameter represents a hybrid metric: it may be affected by both biological and technical variation, as factors such as local light exposure, adaptational state, and/or bleaching by the excitation laser can alter measured response properties.

To quantify how strongly these covariates were reflected in the functional responses, we assessed the accuracy with which each metadata parameter could be predicted from response features (Fig. 5). First, we trained random forest classifiers on the response features (see Methods) to the “chirp” and moving bar stimuli to predict each metadata parameter independently. All metadata parameters were binarised; i.e. age labels were assigned based on postnatal age ( ≤10 weeks vs. >15 weeks), and analogous binary labels were created for all other parameters (see Methods). Training and evaluation of each model were repeated 20 times with different random seeds to ensure robustness. For each run, two constraints were imposed on the train/test split: (1) balanced class representation, and (2) all cells recorded on the same date were held out together to avoid date-specific leakage.

**Fig. 5.**
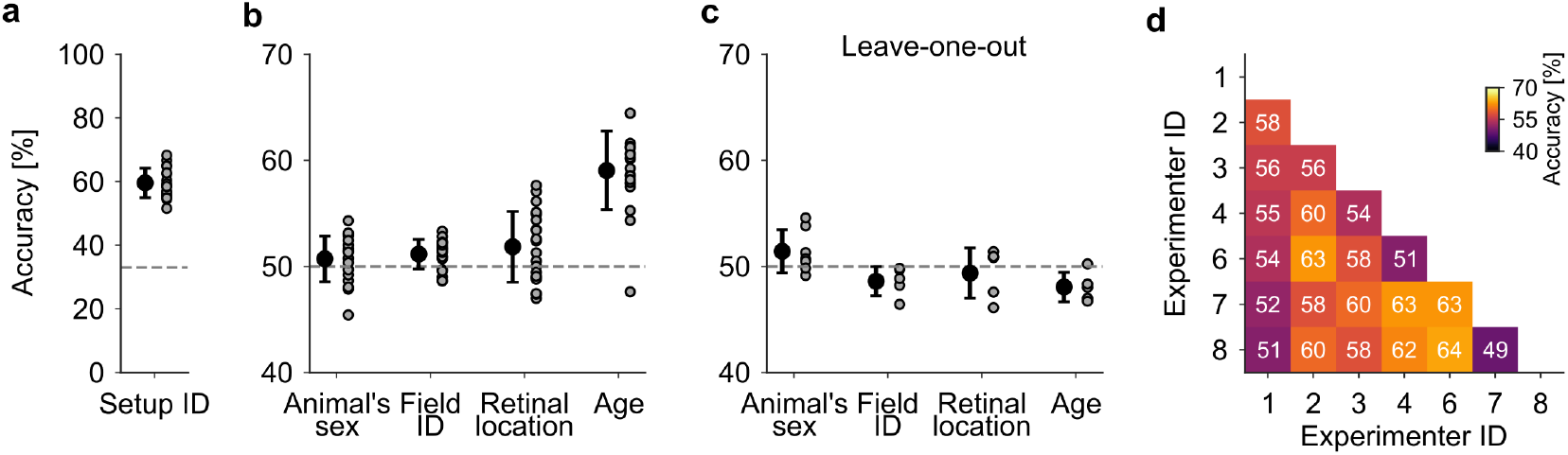
Predictability of experimental and biological variables from GCL cell features. (**a**) Prediction accuracy for the experimental setup ID using a random forest classifier. Each point represents an independent random seed and dataset split. Dashed line indicates chance level (33.3%). (**b**) Prediction accuracy for various metadata categories using a random forest classifier. Each point represents an independent random seed and dataset split. Dashed line indicates chance level (50%). (**c**) Prediction accuracy using a leave-one-out training scheme, where models were trained on all but one experimenter and tested on the held-out one. Error bars represent mean ± SD across dataset splits. (**d**) Cross-experimenter prediction accuracy matrix, showing classification accuracy when discriminating between two experimenters. colour scale indicates accuracy (%, 40–70%).

A classifier was able to distinguish (59.6% ± 4.6%) between different recording setup IDs (Fig. 5a); therefore, for all subsequent classifications, we restricted the data set to the setup ID with the most recordings. Overall, the classifier achieved values around chance level when predicting the animal’s sex, the scan field ID, and the retinal location, indicating that functional response features alone carry little information about these experimental or biological covariates (Fig. 5b). However, it was possible to predict age with 59.0%( ± 3.7%) accuracy, suggesting that age may exert a weak influence on the functional responses. To test whether this age effect reflected biology or experimenter-associated structure, we next applied a leave-one-experimenter-out validation (Fig. 5c). Under this validation scheme, prediction accuracies for scan field ID and retinal location remained close to chance level, while predictability of the animal’s age dropped to chance level. This suggests that the apparent age effect reflected experimenter-specific structure rather than biological differences.

To evaluate experimenter-specific variation, we performed a classifier analysis predicting experimenter identity by training classifiers to predict the experimenter based on the functional recording while restricting data to the same setup (Fig. 5d). Some pairs of experimenters could be distinguished (up to 63% accuracy), while others remained indistinguishable. These results indicate subtle but detectable experimenter-specific effects.

Taken together, accuracies near chance for sex, retinal location, scan field ID, and age (under cross-experimenter validation) indicate that these parameters introduce little systematic structure into the functional responses. In contrast, modestly above-chance accuracies for setup and certain experimenters (≈ 60–63%) reflect weak but detectable technical variation. In the context of high-dimensional neural data, accuracies in this range correspond to minor batch effects that are unlikely to dominate downstream analyses, though adjusting for setup or experimenter may be advisable in sensitive applications. Overall, the All-GCL dataset shows only limited non-biological variability and provides a robust foundation for subsequent analysis.

## Code availability

The DataJoint database was derived from the Python package *djimaging*. The version used for this dataset can be found at github.com/eulerlab/djimaging/releases/tag/all-gcl-v0.1.0. It includes all the code that was used to preprocess the data. The exact schema definition, i.e. which *djimaging* tables have been used and how, and all code for figure generation is available from github.com/eulerlab/all-GCL-manuscript. The GCL classifier is available from github.com/eulerlab/gcl_classifier.

## Data availability

The dataset is found at huggingface.co/datasets/eulerlab/allgcl. The dataset was exported in two formats, (1) a compact *pandas*.*DataF rame* using the .*paquet* format, and (2) detailed single-session exports the Neurodata Without Borders format (27).

## Discussion

The retina has long served as a model system for understanding how neural circuits transform sensory inputs into structured representations of the external world (1, 2). Decades of research have revealed the remarkable diversity of retinal ganglion cell (RGC) types, each encoding specific features of the visual scene such as motion, contrast, or chromaticity (3, 6, 33). Recent advances in functional imaging, single-cell transcriptomics, and connectomics have refined this classification, suggesting 40–50 distinct RGC types in the mouse (7–9). Yet, only a few large-scale functional datasets covering this diversity are publicly available, and even fewer provide standardised stimuli and experimental conditions.

The All-GCL dataset was curated to address this gap. We used a streamlined two-photon Ca^2+^-imaging pipeline and a standardised visual stimulus battery to ensure comparability across preparations collected over nine years. We report extensive metadata that allow users to evaluate biological and technical sources of variability. We further provide functional group labels for all recorded cells by applying a refined GCL classifier, enabling consistent annotations across experiments. Together, these features make the dataset suitable not only for retinal neuroscience but also for benchmarking neural encoding models, testing dimensionality reduction or clustering approaches, and exploring generalisation in biological time-series data. By providing open-access to raw and processed data, we aim to support both hypothesis-driven and data-intensive computational studies.

Compared to previous datasets, which often sampled selected RGC subsets or relied on single preparations, the All-GCL dataset enables large-scale analyses of light-evoked activity across dozens of functional GCL groups. The group-specific topographic maps reveal how distinct GCL types are distributed across the retina, highlighting known organizational principles, such as the ventral enrichment of ‘On’ group cells, and uncovering new, previously under-sampled patterns. Such functional topographies can be directly compared with morphological and transcriptomic atlases, helping to reconcile discrepancies between classification modalities and providing new insights into the spatial specialization of visual processing.

Nevertheless, several limitations must be considered. As the recordings were performed *ex vivo* using Ca^2+^-imaging, they reflect integrated activity rather than precise spike timing, and absolute response latencies may differ from *in vivo* conditions. However, Ca^2+^-dependent signals correlate well with underlying spike trains under similar conditions, providing a reliable proxy for functional classification despite reduced temporal precision (39).

Functional group assignments rely on classifier predictions rather than morphological or molecular ground-truth, and users should account for classification confidence when interpreting results. Although our batch-effect analysis showed only small influences from metadata parameters, some subtle experimenter- or setup-related structure may persist because technical variables are difficult to isolate completely. Additionally, while the dataset covers a large fraction of known GCL groups, types that appear with low sample counts may be genuinely rare or simply infrequent in our recordings relative to their true abundance, and the stimulus repertoire, limited to full-field “chirp” and moving bar paradigms, captures only part of the retina’s functional complexity, which we will address in future releases.

Looking forward, the All-GCL dataset is designed as a “breathing resource”. Future releases will expand its scope with additional mouse lines, including disease models such as *rd10*, and more diverse light stimuli, including colour and naturalistic movies. Together, we expect these efforts to continue transforming the All-GCL dataset into a comprehensive reference for studying the functional organization, diversity, and computation of the mouse retina.

## ACKNOWLEDGEMENTS

We thank Gordon Eske and Merle Harrer for technical support. This work was funded by the German Research Foundation (DFG; CRC 1233, project number 276693517 to PB, TE, KF; GRK 2381, project number 335549539 to TE; Heisenberg Professorship, BE5601/8-1 to PB; Excellence Cluster EXC 2064, project number 390727645 to PB; BE5601/4 to PB; EU 42/10-1, EU 42/12-1 to TE), the Max Planck Society (MPG: M.FE.A. KYBE0004 to KF), the Hertie Foundation (PB), the European Union (ERC,”NextMechMod”, ref. 101039115 to PB), and the the Tistou & Charlotte Kerstan Stiftung (RI-FG P3 EU/SCH 1–2 &3 Dysz to TE, TS). Views and opinions expressed are however those of the authors only and do not necessarily reflect those of the European Union or the European Research Council Executive Agency. Neither the European Union nor the granting authority can be held responsible for them.

## AUTHOR CONTRIBUTIONS

Conceptualisation: DG, JO; Animal protocols: DG, TS; Data acquisition: DG, CC, ND, KPS, FD, TSK, RA, KF; Data curation: JO, DG, FDA, TS; Dataset export: FDA; Formal analysis: DG, JO, TZ; Investigation: JO, DG, FDA, TS; Funding acquisition: TE, PB, KF; Methodology: JO, FDA, ZZ, KF, TE, PB; Resources: ZZ, PB, TE; Software: JO, DG, TZ, FDA, TE; Supervision: TE, PB, KF; Validation: TZ, DG, JO; Visualisation: DG, JO; Writing – original draft: DG, JO, TZ, FDA, TE; Writing – review & editing: DG, JO, TZ, FDA, TE, PB, TS;

## COMPETING FINANCIAL INTERESTS

There are no competing interests to declare.

## Notes

https://huggingface.co/datasets/eulerlab/all-gcl

### Competing Interest Statement

The authors have declared no competing interest.

